# Nuclear speckle protein SON safeguards efficient splicing of GC-rich genes

**DOI:** 10.64898/2026.03.20.713095

**Authors:** Weiyi Fan, Yuenan Zhou, Xinyue Zhang, Chong Tong, Xiaoyu Li, Yafei Yin

## Abstract

Gene architecture in higher eukaryotes exhibits substantial heterogeneity. While most genes follow a canonical pattern of GC-rich exons and AT-rich introns, a subset displays GC-leveled architecture, characterized by uniformly high GC content across both exons and introns. These genes are often associated with nuclear speckles, membraneless compartments enriched in RNA-processing factors, yet the mechanistic basis of this spatial and functional relationship remains unclear. Here, we identify the nuclear speckle protein SON as a critical factor that safeguards the splicing of GC-rich genes. Acute depletion of SON in mouse embryonic stem cells selectively impairs the splicing of short, GC-rich introns. These SON-dependent introns are enriched in highly expressed and functionally essential genes, whose GC-rich architecture contributes to efficient RNA processing and expression. Mechanistically, these introns harbor atypical C-rich, U-poor polypyrimidine tracts at their 3’ splice sites, which exhibit reduced affinity for core splicing factors. SON is recruited to these sites via U2 snRNP and further interacts with SR proteins to stabilize the association of U2 snRNP and U2AFs at these C-rich weak splice sites. Notably, the evolutionary expansion of SON’s intrinsically disordered region is required to promote efficient splicing of GC-rich genes that emerged during evolution. Together, our study suggests that the evolutionary transition toward GC-rich gene architecture enhances gene expression efficiency, with SON acting to safeguard the splicing of this gene class.

## Introduction

GC content varies substantially across eukaryotic genomes and represents a fundamental feature of genome organization.^1^ In higher eukaryotes, genomic regions with distinct GC composition form large domains that differ in gene density, chromatin organization, and transcriptional activity.^2,3^ Genes located in GC-rich regions are often highly expressed and exhibit distinctive structural properties. Gene architecture varies widely across eukaryotic genomes, particularly in the nucleotide composition and length of introns. In most genes, GC-rich exons are flanked by AT-rich introns, a configuration thought to facilitate exon recognition during splicing.^4–7^ However, genomic analyses have revealed a distinct subset of genes that deviate from this canonical architecture and instead display a GC-leveled organization, in which both exons and introns maintain relatively high GC content.^4,5^ These GC-rich genes are typically enriched for short introns, and aberrant splicing of these genes often leads to intron retention, suggesting splicing occur of these genes primarily via intron definition.^4,5^ Despite these observations, the functional significance of GC-rich gene architecture and the mechanisms that ensure efficient RNA processing in such genes remain poorly understood.

Previous studies have noted that GC-rich genes are often positioned in close spatial proximity to nuclear speckles, membraneless nuclear compartments enriched in RNA-processing factors.^8,9^ Nuclear speckles are thought to serve as hubs for RNA metabolism and contain numerous proteins involved in RNA transcription, processing, and export.^10–12^ Consistent with the notion that nuclear speckles facilitate gene expression, genes located near speckles tend to exhibit higher expression levels, and several studies further suggest that nuclear speckles promote efficient splicing of genes located in their vicinity.^8,13–15^ However, the molecular mechanisms underlying this spatial coupling between nuclear speckles and gene splicing remain poorly understood. In particular, whether nuclear speckles, or specific speckle components, directly regulate the splicing of GC-rich genes remains largely unexplored.

Recent work has shown that these structures are largely organized by two scaffold proteins, SON and SRRM2.^11,16^ Interestingly, both proteins contain large intrinsically disordered regions (IDRs) that have undergone substantial expansion during evolution in higher eukaryotes. Functional studies of these proteins have suggested their involvement in RNA processing.^17–19^ In particular, depletion of SON results in widespread splicing defects affecting a subset of transcripts, especially those containing weak splice sites.^18^ These findings suggest that SON acts as an auxiliary splicing regulator that facilitates efficient spliceosome assembly or stabilization on challenging pre-mRNA substrates. In addition, SON has been implicated in the proper expression of genes involved in cell cycle progression, pluripotency maintenance, and mutations in SON cause intellectual disability syndrome, underscoring its importance in gene regulatory programs and human disease.^18–22^ However, the molecular mechanisms by which SON regulates splicing, as well as the specific classes of genes or introns that depend on SON for efficient processing, remain poorly understood.

Here, we show that the nuclear speckle protein SON plays a critical role in safeguarding the efficient splicing of GC-rich genes. Using rapid SON degradation in mouse embryonic stem cells combined with nascent transcript sequencing and quantitative splicing analysis, we find that SON depletion causes widespread splicing defects that predominantly affect short, GC-rich introns containing C-rich, U-poor 3’ splice sites. Mechanistically, SON associates with the U2 snRNP complex and stabilizes spliceosome assembly at these weak splice sites, thereby promoting efficient splicing. These findings reveal a mechanism that supports efficient RNA processing of GC-rich genes and suggest that SON may have evolved to safeguard this distinctive gene architecture in higher eukaryotes.

## Results

To investigate the direct function of SON, we generated a degron mouse embryonic stem cell (mESC) line enabling rapid depletion of SON (SON^dTAG^).^23^ In this system, treatment with dTAG efficiently degrades the majority of SON protein within 2 hours (Fig. 1A). Notably, prolonged SON degradation severely impaired mESC survival. Consistent with this phenotype, the stress-responsive apoptotic protein p53 accumulated substantially after 8 hours of SON depletion (Fig. 1A, B). Consistent with previous observations, depletion of SON also caused marked alterations in nuclear speckle morphology, resulting in fewer but enlarged speckles (Fig. 1C).^16,18^ To directly assess the impact of SON depletion on RNA processing and RNA localization, we labeled newly synthesized RNA with 4-thiouridine (4sU) following 3 hours of SON degradation. Labeled transcripts were subsequently purified and analyzed by sequencing from multiple RNA pools, including total RNA (4sU-seq), polyadenylated RNA (4sU-pA-seq), transient transcriptome sequencing (TT-seq), and subcellular RNA fractions (sub-4sU-seq) (Fig. 1D).^24–26^ These approaches allowed us to capture nascent transcription and RNA processing events with high temporal resolution following SON depletion.

**Figure 1.**
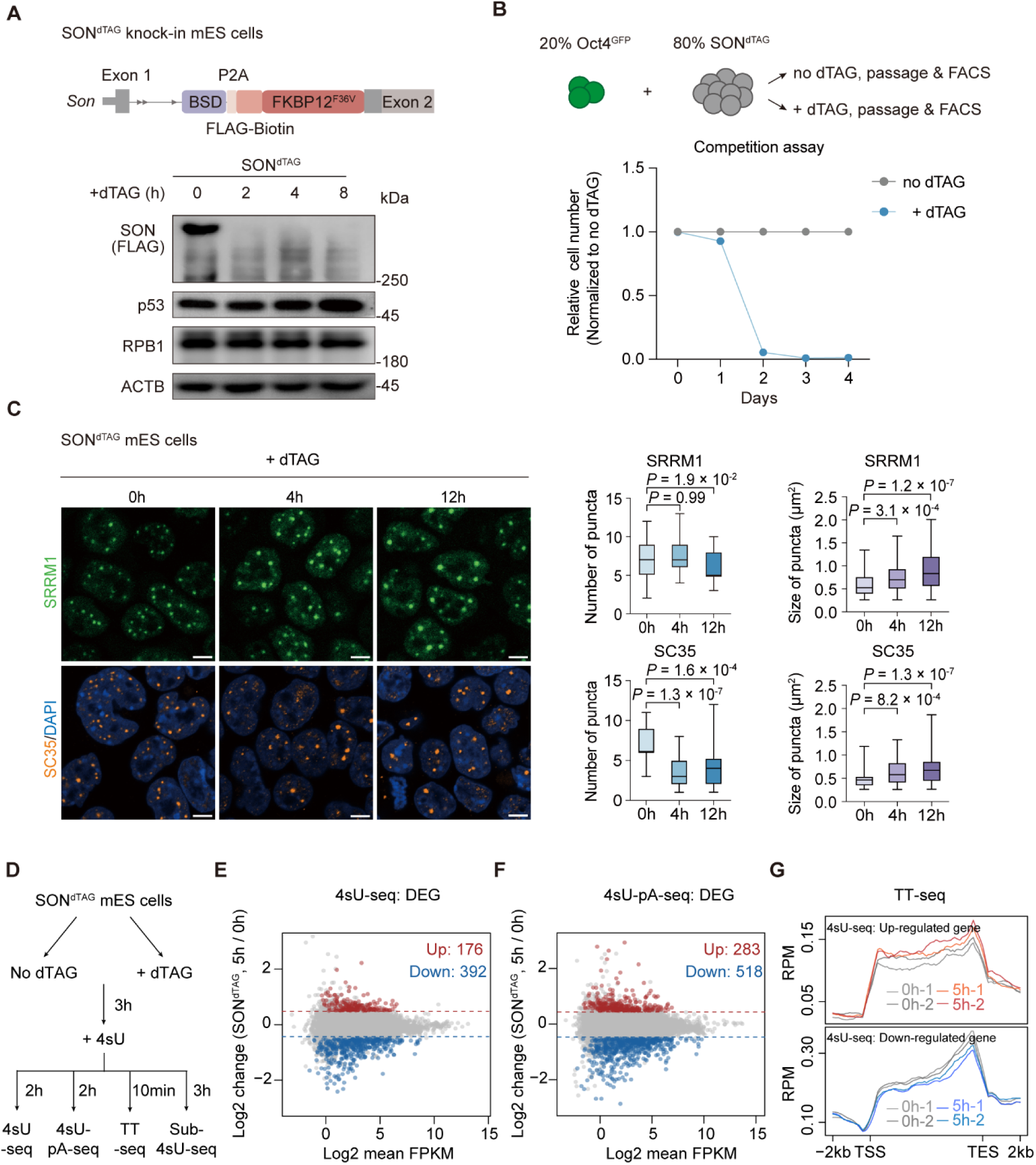
Rapid depletion of SON induces growth defects, alters nuclear speckle organization and has minimal effects on transcription. (A) Top, schematic of the dTAG system used for rapid depletion of SON protein. The endogenous SON locus was tagged with FKBP12^F36V^ at exon 2. Bottom, Western blot analysis shows time-dependent SON degradation after dTAG treatment (0, 2, 4, and 8 h), along with the corresponding protein levels of p53 and RPB1. (B) Top, schematic of the competition assay to examine the proportion of SON^dTAG^ mESCs under long-term dTAG treatment. Bottom, relative abundance of dTAG-treated cells compared with untreated controls across passages. Lines represent mean values from two independent experiments. Flow cytometry gating strategies are shown in the Methods. (C) Maximum intensity projection of images of SON^dTAG^ mESCs following dTAG treatment for the indicated times (0, 4, and 12 h). Top, quantification of SRRM1-GFP puncta from live-cell imaging. Puncta number represents the number of puncta per cell (0 h, *n =* 31 cells; 4 h, *n =* 23 cells; 12 h, *n =* 26 cells). Puncta size represents individual puncta measurements (0 h, *n =* 109 puncta; 4 h, *n =* 107 puncta; 12 h, *n =* 105 puncta). Bottom, quantification of SC35 puncta by immunofluorescence. Puncta number represents the number of puncta per cell (0 h, *n =* 22 cells; 4 h, *n =* 24 cells; 12 h, *n =* 26 cells). Puncta size represents individual puncta measurements (0 h, *n =* 95 puncta; 4 h, *n =* 34 puncta; 12 h, *n =* 76 puncta). Only puncta with an area > 0.25 μm² were included in the size analysis. Nuclei were stained with DAPI. *P* values, two-sided *t*-tests. Scale bar, 5 μm. (D) Schematic of the 4sU-seq, 4sU-pA-seq, TT-seq and subcellular fraction 4sU-seq (Sub-4sU-seq) of SON^dTAG^ mESCs with or without dTAG treatment. (E) MA plot showing the upregulated (red, log2FC > 0.5, *P* < 0.05) and downregulated (blue, log2FC < -0.5, *P* < 0.05) genes identified by 4sU-seq of the acute degradation of SON. *P* value*s*, two-sided *t*-test. (F) MA plot showing the upregulated (red, log2FC > 0.5, *P* < 0.05) and downregulated (blue, log2FC < -0.5, *P* < 0.05) genes identified by 4sU-pA-seq of the acute degradation of SON. *P* value*s*, two-sided *t*-test. (G) Metaplot of genes upregulated or downregulated in 4sU-seq, showing minimal changes in TT-seq signal.

Acute depletion of SON resulted in dysregulation of only a small number of genes in both 4sU-seq and 4sU-pA-seq, and TT-seq showed that transcription of these genes was not substantially altered, suggesting that SON does not directly regulate gene transcription (Fig. 1E-G). We therefore examined the impact of SON depletion on pre-mRNA splicing by quantifying 3’ splice site (3’ss) usage for each intron (Fig. 2A).^24^ This analysis identified 3,245 introns with significantly decreased splicing and only 185 introns with increased splicing following SON depletion in 4sU-seq (|log2FC| > 1, *P* < 0.05). A similar pattern was observed in the 4sU-pA-seq dataset (Fig. 2A and S1A). We therefore defined introns exhibiting significantly reduced splicing as SON-regulated introns, whereas introns showing minimal splicing change (−0.1 < log2FC < 0.1) were defined as unregulated controls.

**Figure 2.**
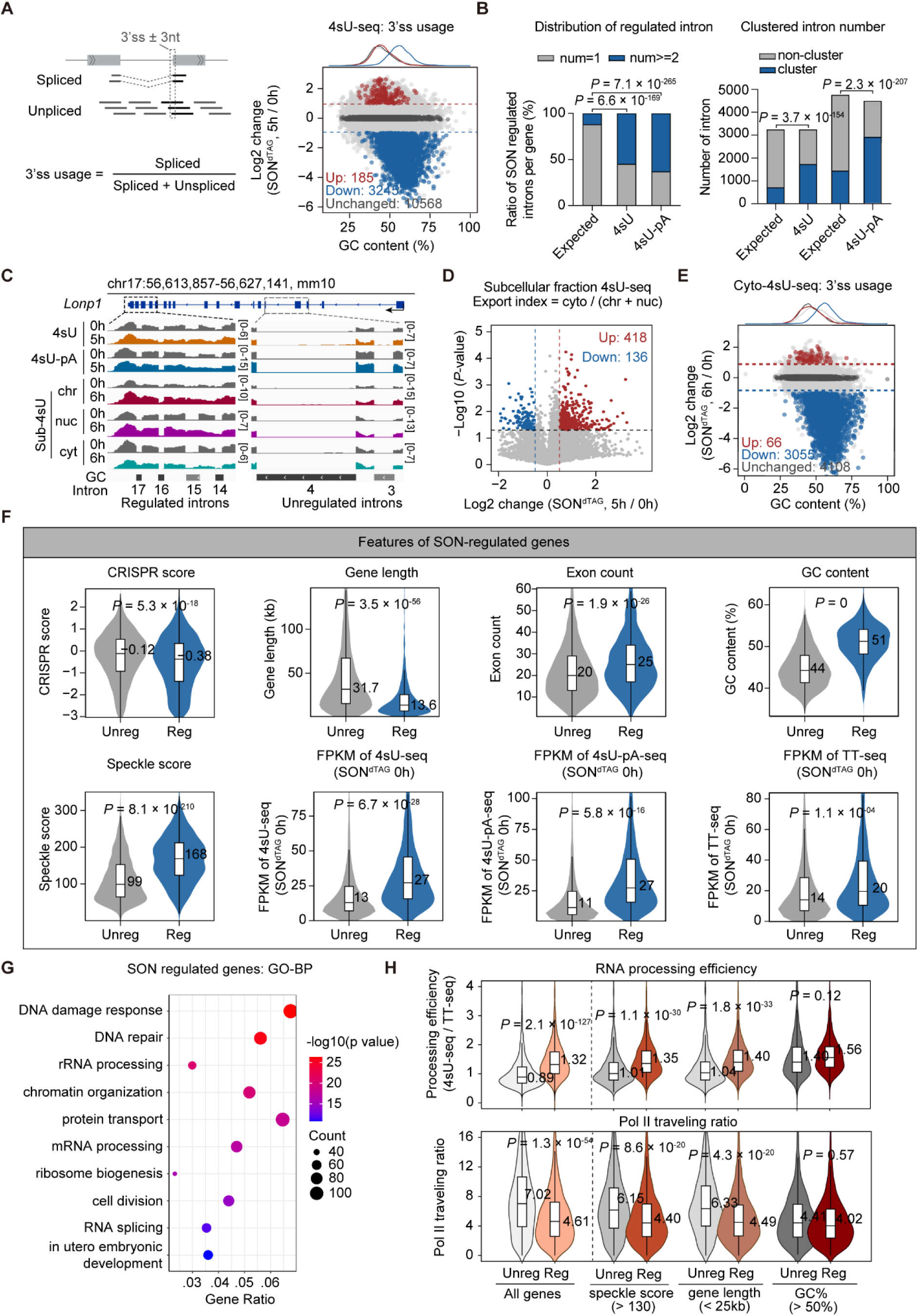
Feature analysis of SON-regulated genes. (A) Left, Schematic of splicing efficiency analysis. Reads overlapping with the region used to calculate the spliced/unspliced ratio are indicated by a dashed box. Right, analysis of 3’ss usage in SON^dTAG^ mESCs by 4sU-seq reveals that SON-regulated introns have high GC content. (B) Comparison of the proportion of SON-regulated introns per gene and intron clustering number between 4sU-seq and 4sU-pA-seq datasets. Expected values were estimated by random sampling of an equal number of introns, followed by calculation of the fraction occurring within the same gene and forming clusters. Statistical significance between conditions was assessed using Fisher’s exact test. (C) IGV track shows increased intron retention in the *Lonp1* gene upon SON depletion, with introns 14-17 forming a cluster of SON-regulated introns. Introns were grouped by GC content into three categories (< 44, 44–50, ≥ 50%), with darker shades of gray corresponding to higher GC content. (D) Volcano plot showing changes in RNA transport efficiency upon SON depletion. A total of 136 genes exhibited a slight decrease, whereas 418 genes showed a significant increase in RNA transport efficiency. (E) 3’ss usage analysis in SON^dTAG^ mESCs by cyto-4sU-seq reveals that SON-regulated introns have high GC content. (F) Boxplots show gene features of host genes harboring SON-regulated introns, defined by integrating 4sU-seq and 4sU-pA-seq datasets. (G) GO analysis of biological processes for the gene set shown in (F). (H) Top, SON-regulated genes (Reg) show higher RNA processing efficiency (4sU-seq FPKM / TT-seq FPKM) than unregulated genes (Unreg). All genes: Unreg, *n =* 5,798; Reg, *n =* 1,714. Subsequent analyses were restricted to expressed genes (mean 4sU-seq FPKM > 10) and further stratified by nuclear speckle score (>130; Unreg, *n =* 1,193; Reg, *n =* 1,060), gene length (<25 kb; Unreg, *n =* 1,532; Reg, *n =* 1,125), or GC content (>50%; Unreg, *n =* 479; Reg, *n =* 897). Bottom, SON-regulated genes exhibit higher RNAPII traveling ratios (see Methods). Gene sets are as in the top panel, with additional filtering for genes with maximum Pol II read count >50 in the gene body. Final gene numbers (left to right): 5,656, 1,649, 1,171, 1,026, 1,475, 1,076, 453, and 860. *P* value*s*, two-sided *t*-tests.

Strikingly, SON-regulated introns displayed a strong gene distribution bias, with 3,245 and 4,759 regulated introns located within 1,161 (4sU-seq) and 1,346 (4sU-pA-seq) genes—significantly fewer genes than expected by chance (*P* < 6.6×10^-169^, Fig. 2B). Moreover, inspection of individual genes and global analysis further revealed that SON-regulated introns frequently appeared in clusters, with two or more regulated introns positioned adjacent to each other, implying coordinated or synergistic regulation of splicing within local intron groups (Fig. 2B, C, and S1B-D). In addition, depletion of SON had limited impact on global RNA export, while many SON-regulated introns remained unspliced even after export to the cytoplasm, indicating that depletion of SON abolishes, rather than merely delays, the splicing of these introns (Fig. 2D, E, and S1B-D).

We defined genes containing introns with reduced splicing in either 4sU-seq or 4sU-pA-seq as SON-regulated genes. Feature analysis comparing these genes with unregulated controls revealed several distinctive properties: SON-regulated genes tend to be GC-rich, shorter in length, and located closer to nuclear speckles, resembling features previously described for genes exhibiting GC-leveled architecture (Fig. 2F).^8,13^ In addition, these genes show higher essentiality scores, and gene ontology analysis indicates enrichment for processes such as DNA damage response, RNA processing, and chromatin organization, suggesting that SON-regulated genes are largely associated with housekeeping functions (Fig. 2F, G). Notably, compared with unregulated controls, SON-regulated genes exhibited higher expression levels in both stable and nascent RNA, as well as substantially higher RNA processing efficiency (measured as the 4sU-seq/TT-seq ratio) and elongated RNA Polymerase II (RNAPII) ratio (measured by travel ratio of RNAPII) (Fig. 2F, H).^27^ To determine which feature primarily contributes to the faster processing of SON-regulated genes, we restricted the analysis to robustly expressed genes (FPKM > 10 in 4sU-seq) and compared processing efficiencies under different feature constraints. Strikingly, controlling for GC content, but not gene length or nuclear speckle proximity, largely eliminated the difference in RNA processing efficiency and elongated RNAPII ratio between SON-regulated and unregulated genes (Fig. 2H). These results suggest that the high GC content of SON-regulated genes is a major factor facilitating their processing and higher expression, while SON functions to safeguard efficient splicing within this gene class.

Consistent with the properties of genes exhibiting a GC-leveled architecture, SON-regulated introns were shorter, with higher GC content compared with unregulated controls (Fig. 3A).^4^ Intriguingly, these introns were less conserved among vertebrates, both at the level of the full intron sequence and at splice sites, indicating that SON-regulated introns may evolve more rapidly (Fig. 3A). Motif analysis of key splicing regulatory elements revealed marked sequence differences between SON-regulated and unregulated introns, particularly at the 3’ss. In canonical splice sites, the polypyrimidine tract (PPT) is typically T-rich (U-rich in RNA). In contrast, SON-regulated introns displayed C-rich PPT sequences (Fig. 3B). Consistent with this observation, analysis of alternative splicing events using rMATS indicated that exon skipping and intron retention were the most frequent events upon depletion of SON, and motif analysis showed that the 3’ss of affected introns in both categories were similarly C-rich (Fig. S2).

**Figure 3.**
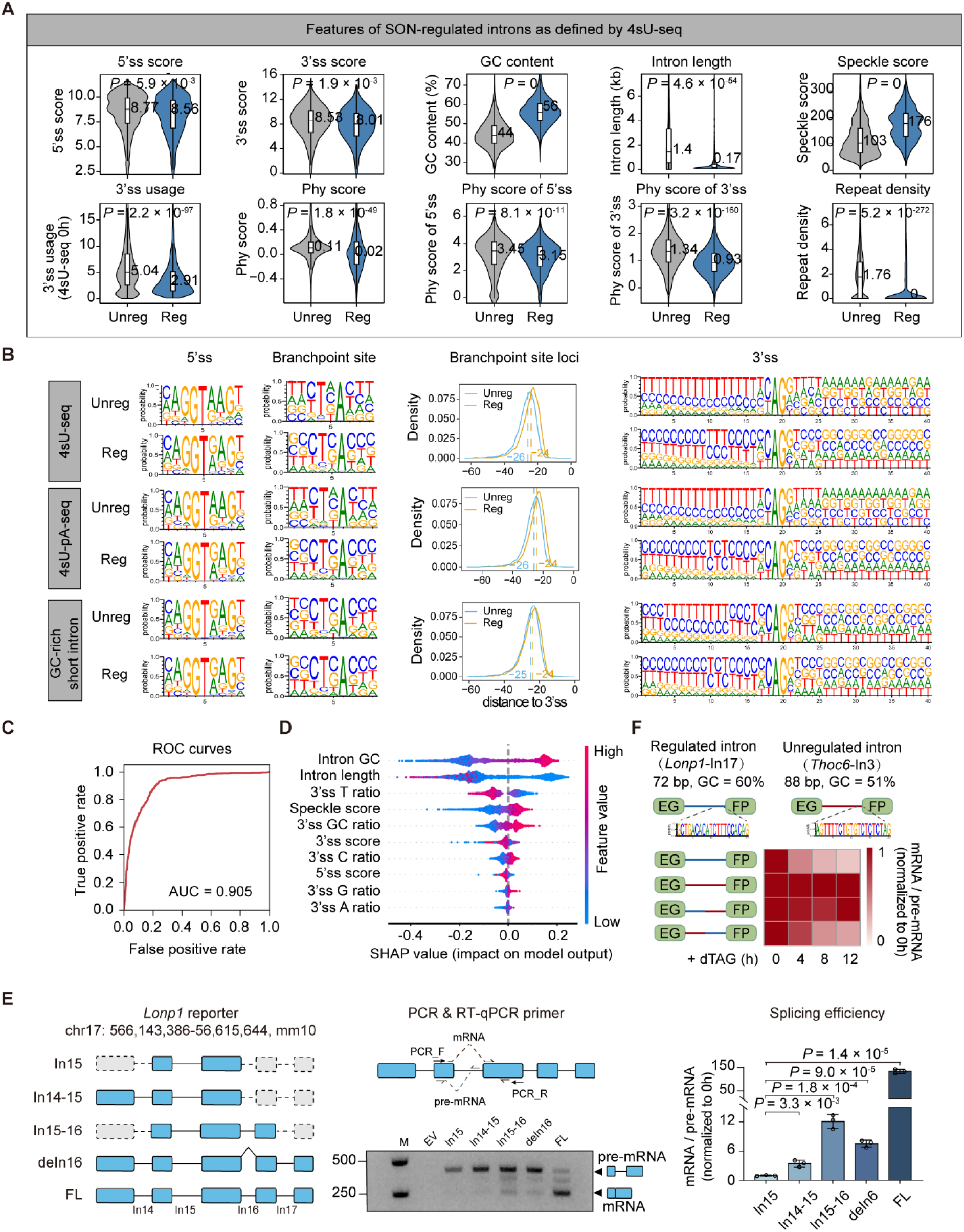
SON promotes efficient splicing of GC-rich genes. (A) Boxplots show features of SON-regulated introns identified by 4sU-seq. Phylogenetic conservation scores were obtained using phyloP based on multi-species genome alignments. (B) Sequence features of SON-regulated and unregulated introns were analyzed using 4sU-seq (Unreg: 10,568; Reg: 3,245), 4sU-pA-seq (Unreg: 11,860; Reg: 4,759), and GC-rich short introns (Unreg: 5,520; Reg: 5,965; see Methods). Features shown include 5’ss motifs, branchpoint site motifs, branchpoint-to-3’ss distance distributions, and 3’ss motif. SON-regulated introns were defined as GC-rich short introns (GC% > 50) with a maximum spliced or unspliced read count > 10, *P* < 0.2, and log2 fold change < 0 (*n =* 5,965). SON-unregulated introns were defined as GC-rich short introns (GC% > 50) with maximum spliced or unspliced read count > 10, subdivided into two groups: introns with log2 fold change > 0 and *P* < 0.2 (*n =* 395), and introns with *P* ≥ 0.2 (*n =* 5,520), giving a total of 5,915 unregulated introns. (C) Receiver operating characteristic (ROC) curve of the predictive model built using features of SON-regulated introns, showing an area under the curve (AUC) of 0.905. (D) SHAP analysis identified features that significantly promote SON-mediated intron regulation. (E) Left, schematic diagrams of each reporter transcript (exons, blue boxes; single intron, black). Dashed lines indicate regions not included in the reporter. Middle, positions of PCR and RT–qPCR primers are indicated, along with splicing efficiency of each reporter assessed by PCR analysis. Right, RT–qPCR analysis of splicing efficiency (mRNA / pre-mRNA) for each reporter. Data are shown as mean ± s.d. (*n* = 3 independent replicates). (F) Top, schematics of EGFP minigene reporters containing an EGFP coding sequence interrupted by either a SON-regulated intron (*Lonp1*-In17) or unregulated intron (*Thoc6*-In3). Bottom, heatmaps showing RT–qPCR analysis of the splicing efficiency (mRNA/pre-mRNA) for the indicated EGFP minigene reporters (structures shown at left) upon SON^dTAG^ degradation. The reporters contain four different introns, and detailed schematics and descriptions are shown in Fig. S3E. *P* value*s*, two-sided *t*-tests.

To quantitatively evaluate the contribution of different intron features to SON regulation, we constructed a machine-learning model using sequence and genomic features of regulated and unregulated introns (Fig. 3C, D). Feature importance analysis revealed that intron GC content, intron length, speckle proximity, and particularly 3’ss sequence composition were the strongest predictors of SON dependency. Inspection of individual loci further showed that, for most regulated genes, only a subset of their introns is affected by SON depletion, and some SON-regulated genes are located relatively far from nuclear speckles, suggesting that spatial positioning is not the primary determinant of SON regulation (Fig. 2C and S1C, D). To further refine the sequence characteristics associated with SON regulation, we performed motif analysis restricted to high-GC short introns.^4^ Importantly, the C-rich, T-poor 3’ss motif remained strongly enriched in SON-regulated introns under this condition, where most other sequence features were comparable between regulated and unregulated groups (Fig. 3B). These results indicate that 3’ss sequence composition is a key determinant of SON-dependent regulation.

To further test this idea, we expressed SON-regulated introns exogenously as splicing reporters. Both single regulated introns and clustered intron reporters exhibited impaired splicing following acute SON depletion, indicating that SON-dependent regulation is largely independent of chromatin environment (Fig. S3). Notably, clustered intron reporters showed higher splicing efficiency than single-intron reporters, supporting synergistic splicing effect among adjacent regulated introns (Fig. 3E). Importantly, exchanging the 3’ portion, but not the 5’ portion, of a SON-regulated intron with that of an unregulated GC-rich intron abolished SON dependency (Fig. 3F and S3E). Together, these results strongly demonstrate that 3’ss sequence features are the primary determinants of SON-mediated regulation.

Previous studies have shown that reduced uridine content in the polypyrimidine tract weakens the affinity of U2AF2 for the 3’ss and lowers splicing efficiency.^28,29^ Consistent with this, SON-regulated introns exhibited substantially lower splicing efficiency than unregulated controls (Fig. 3A). We therefore hypothesized that SON promotes splicing by stabilizing the interaction of U2 snRNP and U2AF factors with the 3’ss of regulated introns. Indeed, acute SON depletion caused a much stronger reduction in U2 snRNP and U2AF2 binding at the 3’ss of SON-regulated introns compared with unregulated introns (Fig. 4A). Proteomic analysis using TurboID further revealed that the SON interactome is highly enriched for splicing regulators, particularly components of the U2 snRNP (Fig. 4B-D).^30^ To further probe this interaction, we treated cells with Pladienolide B, which targets SF3B1 and disrupts the interaction between U2 snRNP and pre-mRNA.^31^ PladB treatment reduced SON interactions with SR proteins and factors involved in later stages of splicing, such as SNRNP200, whereas interactions with most U2 snRNP components remained largely intact, with the exception of SF3B3 (Fig. 4B, D, E). In addition, PladB treatment reduced SON association with pre-mRNAs from both regulated and unregulated genes, suggesting that SON associates with nascent transcripts largely through its interaction with the U2 snRNP complex (Fig. 4F). Together, these results support a model in which SON targets pre-mRNAs via U2 snRNP and associated splicing factors, further interacting with SR proteins and stabilizing the recruitment of U2 snRNP and U2AFs at C-rich 3’ss, thereby promoting efficient splicing of GC-rich introns within the genes it regulates.

**Figure 4.**
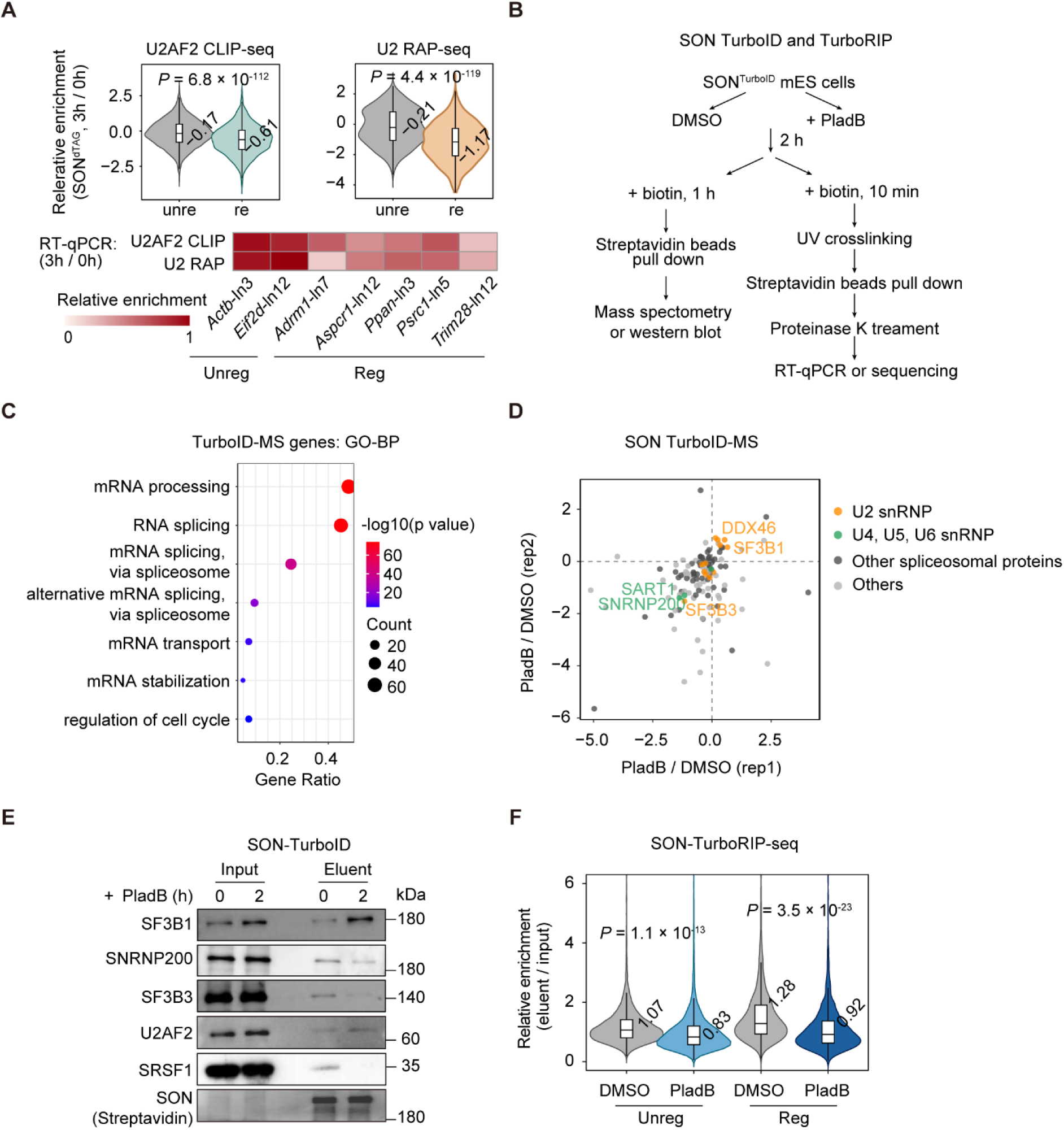
SON stabilizes the recruitment of U2 snRNP and U2AFs to promote efficient splicing of GC-rich introns. (A) U2AF2 CLIP and U2 RAP-seq show that SON depletion reduces binding of these factors to SON-regulated introns. Heatmap of RT-qPCR-based enrichment of U2AF2 and U2 RAP on SON-regulated and unregulated introns following SON depletion is shown below. (B) Schematic of the TurboID and TurboRIP of SON^TurboID^ mESCs with DMSO or PladB treatment. (C) GO analysis of biological processes for the TurboID-MS enriched genes (*n =* 126). (D) Scatter plot showing the changes of the abundance of proteins captured by SON TurboID after DMSO or Pladienolide B (PladB) treatment in SON^TurboID^ mESCs. (E) Western blot analysis of SON TurboID captured proximal proteins in SON^TurboID^ mESCs with DMSO or PladB treatment. (F) TurboRIP-seq analysis shows that PladB treatment markedly reduces SON association with SON-regulated (*n =* 1,716) and unregulated genes (*n =* 5,787). *P* value*s*, two-sided *t*-tests.

Mammalian SON contains a conserved structured C-terminal region, a less conserved disordered RS domain, and an N-terminal IDR that has undergone substantial expansion during evolution (Fig. 5A and S4A).^16,18^ To assess the contribution of these domains, we generated a series of truncation constructs and evaluated their ability to rescue SON depletion. Surprisingly, deletion of the conserved C-terminal region, but not constructs lacking the less conserved RS domain or portions of the N-terminal IDR, partially restored cell survival and the splicing defects of analyzed SON-regulated introns (Fig. 5A, B). Notably, the rescue efficiency correlated with the degree of SON colocalization with nuclear speckles (*R* = 0.86, *P* = 4.9 × 10^-6^), suggesting that proper SON localization to nuclear speckles is closely linked to its splicing regulatory function (Fig. 5A-E).

**Figure 5.**
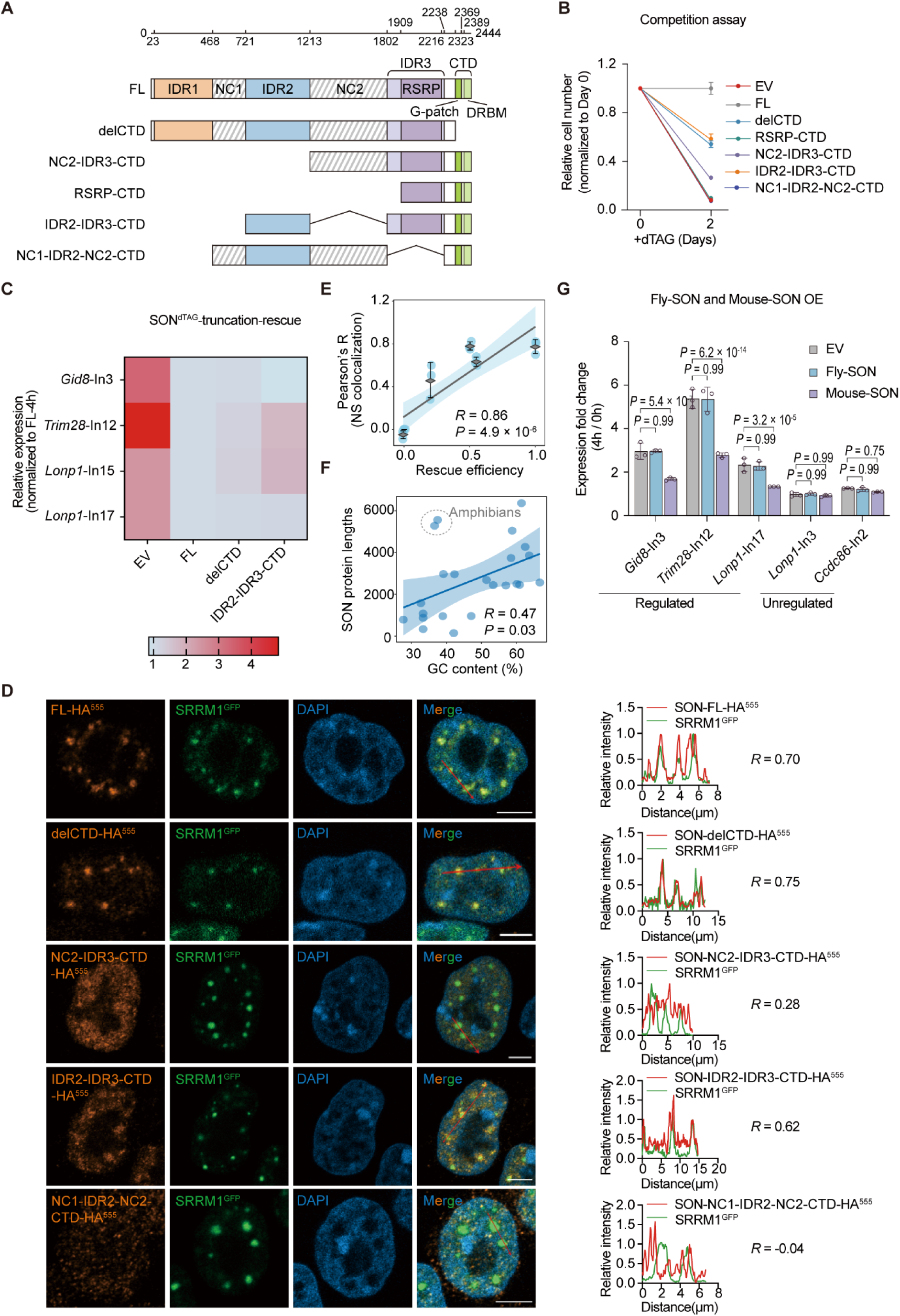
Evolutionary expansion of SON’s IDR is required for its proper function in mES cells. (A) Schematic representation of SON truncation constructs based on the domain architecture of SON. The scale above indicates the amino acid positions along the SON protein. FL, full-length; IDR, intrinsically disordered region; NC, negatively charged region; CTD, C-terminal domain; RSRP, arginine- and serine-rich region; DRBM, double-stranded RNA binding motif. (B) Relative cell numbers of SON^dTAG^ mESCs rescued with different SON truncation constructs following long-term dTAG treatment (2 days). EV, empty vector. FL, full length of *Son*. Cell numbers were normalized to Day 0. Lines represent mean values from three independent experiments. (C) Heatmap showing RT–qPCR analysis of relative pre-mRNA levels of SON-regulated introns (*Gid8*-In3, *Trim28*-In12, *Lonp1*-In15, *Lonp1*-In17) in SON^dTAG^ mESCs rescued with different SON truncation constructs following 4 h dTAG treatment. Expression levels were normalized to *Actb* exon (*Actb*-E) and are shown relative to SON-FL. (D) Representative immunofluorescence images showing the colocalization of SON truncation constructs with SRRM1^GFP^ in mESCs. DAPI marks the nucleus. Line-scan analysis of fluorescence intensity along the red line in the merged images is shown on the right, illustrating the spatial correlation between SON constructs and SRRM1. Pearson correlation coefficients (*R*) are indicated. Scale bar, 5 μm. (E) The rescue efficiency of SON truncations correlates with their degree of nuclear speckle co-localization, as determined by immunofluorescence (Fig. 5C, D). Pearson correlation coefficient (*R*) and corresponding *P* value*s* are shown. (F) SON protein length is correlated with the median GC content of short introns, although some amphibian species show deviations from this trend. GC content of short introns in different species is shown in Fig. S4A. Pearson correlation coefficient (*R*) and corresponding *P* value*s* are shown. (G) RT–qPCR analysis of pre-mRNA levels of SON-regulated introns (*Gid8*-In3, *Trim28*-In12, *Lonp1*-In17) and unregulated introns (*Lonp1*-In3, *Ccdc86*-In2) in SON^dTAG^ mESCs overexpressing either Fly SON or Mouse SON, following dTAG treatment at 0 h and 4 h. Expression levels were normalized to *Actb*-E and are shown as fold change relative to the 0 h condition (4 h / 0 h). Data are shown as mean ± s.d. (*n* = 3 independent replicates). Statistical significance was determined using two-way ANOVA.

The evolutionary expansion and functional importance of SON’s IDR, together with the rapidly evolving and distinctive features of SON-regulated GC-rich introns, raise the possibility that these two elements have coevolved. Comparative evolutionary analyses revealed a positive correlation between the expansion of the SON’s IDR and the emergence of short, GC-rich introns across species (*R* = 0.47, *P* = 0.03) (Fig. 5F and S4A, B). Consistent with this notion, expression of *Drosophila* SON failed to rescue the splicing deficiency caused by depletion of mouse SON (Fig. 5G). Together, these observations suggest that the evolutionary expansion of SON’s IDR contributes to safeguarding the efficient splicing of GC-rich introns that emerged during evolution (Fig. 6).

**Figure 6.**
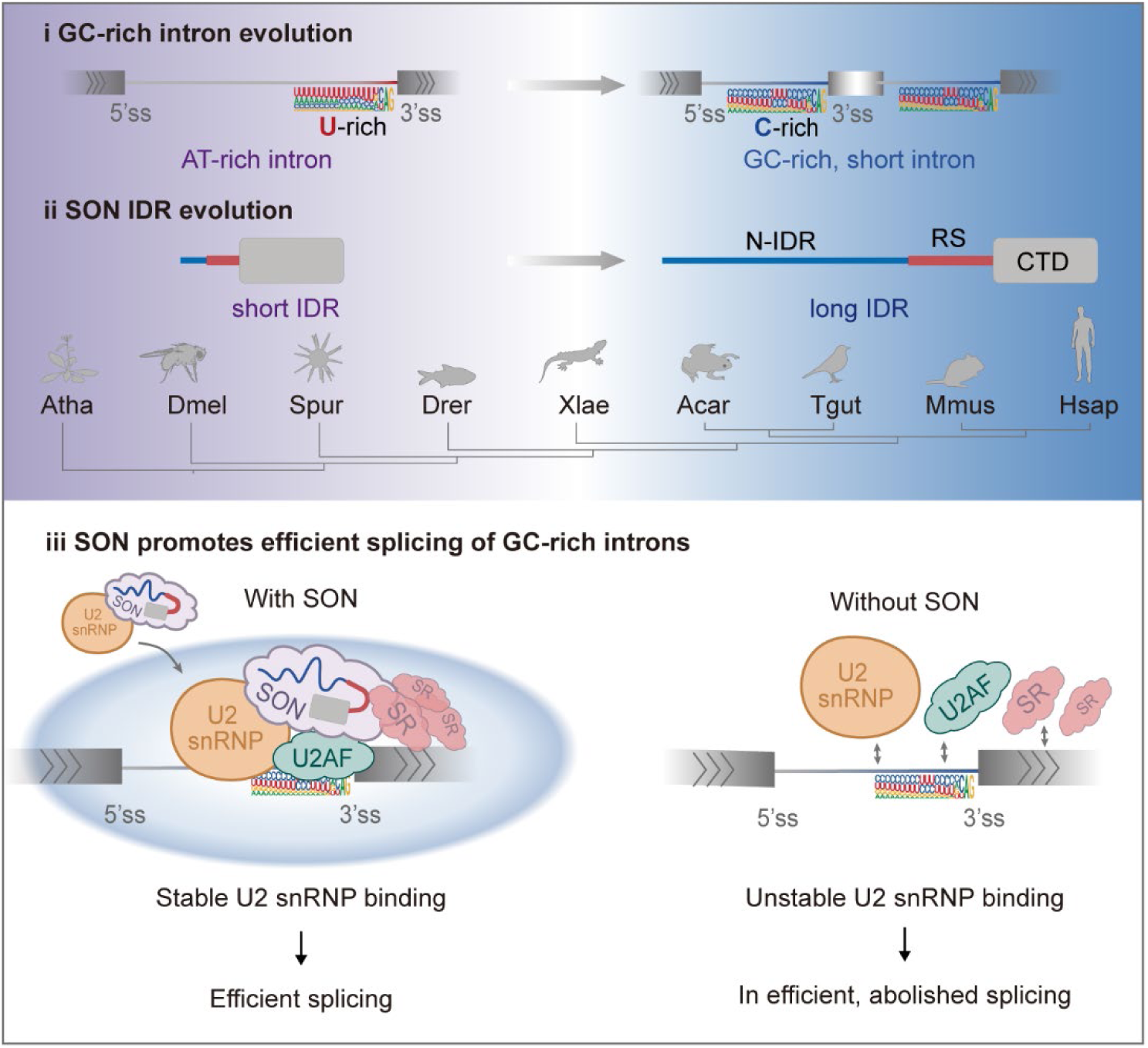
Model linking the evolutionary expansion of SON’s IDR to the emergence and regulation of short GC-rich introns. (i) Intron architecture shifts from predominantly long introns to shorter, GC-rich introns, accompanied by a change in 3’ss composition from U-rich to C-rich sequences. (ii) The N-terminal IDR of SON progressively expands during evolution, and the emergence of short GC-rich introns across species is shown below. The phylogenetic tree was obtained from TimeTree5. Atha: *Arabidopsis thaliana*, Dmel: *Drosophila melanogaster*, Spur: *Strongylocentrotus purpuratus*, Drer: *Danio rerio*, Xlae: *Xenopus laevis*, Acar: *Anolis carolinensis*, Tgut: *Taeniopygia guttata*, Mmus: *Mus musculus*, and Hsap: *Homo sapiens*. (iii) In the presence of SON, SON is recruited to short GC-rich introns via interaction with U2 snRNP, where it interacts with SR proteins, potentially through its RS domain, and stabilizes binding of U2 snRNP and U2AFs at weak C-rich 3’ss. The expanded IDR of SON may further promote formation of condensates that concentrate spliceosomal components and facilitate synergistic splicing. In the absence of SON, binding of U2 snRNP and U2AFs at these sites is unstable, resulting in inefficient or abolished splicing of these introns.

In summary, our study identifies SON as a key regulator that promotes efficient splicing of GC-rich introns by stabilizing spliceosome assembly at C-rich 3’ss. Our findings further suggest that the evolutionary transition toward GC-rich gene architecture enhances RNA processing and gene expression efficiency, with SON acting to safeguard the splicing of this gene class.

## Discussion

Gene architecture is highly heterogeneous across eukaryotic genomes, with GC-rich gene architectures preferentially enriched within megabase-scale GC-rich regions, known as isochores.^1^ Previous studies have shown that these regions are gene-dense, positioned in close proximity to nuclear speckles, and associated with high levels of gene expression, However, their functional significance and underlying mechanism of this arrangement remains incompletely understood.^8,15^ Our results suggest that the increase in GC-content during evolution of these genes may facilitate their faster processing and efficient expression (Fig. 2H). Specifically, although SON-regulated introns exhibit intrinsically lower splicing efficiency, their organization into short introns and clustered arrangements may enable synergistic splicing, thereby compensating at the gene level. In addition, the spatial proximity of these genes, both to each other and to nuclear speckles enriched in transcriptional and RNA processing factors, may further promote coordinated and efficient gene regulation.

Intriguingly, our analysis also shows that GC-rich genes exhibit a lower RNAPII travel ratio, which is often interpreted as more efficient release from promoter-proximal pausing (Fig. 2H).^27,32^ This is supported by previous in vitro studies suggesting that co-transcriptional folding of GC-rich nascent RNA may impose an energy barrier to pausing by impeding RNAPII backtracking.^33^ However, it is also reported that GC content within gene bodies is negatively correlated with transcription elongation speed, which may contribute to the reduced travel ratio we observed.^34,35^ Under such conditions, slower transcription elongation could provide a larger temporal window for co-transcriptional splicing, thereby facilitating the processing of these intrinsically less efficient, SON-dependent introns.

Nevertheless, these features suggest that GC-rich gene architecture may represent an evolutionary strategy that supports high and efficient gene expression in complex eukaryotic genomes. However, the distinctive sequence composition of GC-rich introns also imposes constraints on splicing, particularly through the emergence of C-rich polypyrimidine tracts that weaken canonical splice-site recognition. Our findings indicate that SON functions, with a particular dependence on its expanded IDR region, to overcome these limitations by stabilizing spliceosome assembly at such noncanonical splice sites. While we do not exclude the possibility that the evolutionary expansion of SON’s IDR reduces the dependence of splicing on strict sequence features, thereby relaxing selective constraints and facilitating the transition toward GC-rich introns (Fig. 6, panel i and ii), this possibility remains to be further investigated.

Our study suggests that 3’ss sequence features are a major determinant of SON-dependent regulation. In particular, SON-regulated introns are characterized by U-poor, C-rich 3’ss, which exhibit weak U2AF2 binding affinity and reduced spliceosome assembly efficiency in vitro.^28,29^ This is also supported by previous study showing that SON preferentially regulates introns with weak splice sites.^18^ Our results further indicate that this specificity is not driven by selective targeting of SON to particular introns, as inhibition of splicing abolishes SON association with both regulated and unregulated introns (Fig. 4F). Instead, the specificity appears to be determined by the intrinsic properties of these C-rich 3’ss, leading to more dynamic and less stable binding of U2AF and U2 snRNP. In addition, GC-rich exon sequences, which are relatively depleted of GA-rich motifs that serve as major binding sites for SR proteins (Fig. 3B), may further contribute to reduced and dynamic SR protein binding.^36,37^ Thus, our study supports a model in which SON is recruited to pre-mRNAs via its interaction with U2 snRNP and functions as a scaffold that further interacts with SR proteins, potentially mediated by its RS domain, as well as other splicing regulators. Through reinforcing multiple unstable and dynamic interactions, SON promotes efficient spliceosome assembly and splicing at these weak 3’ss (Fig. 6iii).

While our results support a central role for SON in promoting the splicing of GC-rich genes, the precise relationship between SON’s molecular function and its role in nuclear speckle organization remains to be fully resolved. Although acute depletion of SON leads to significant splicing defects in GC-rich introns, nuclear speckles are not abolished under these conditions (Fig. 1C), suggesting the existence of nuclear speckle structure is not sufficient to support efficient splicing of GC-rich genes. At the same time, our rescue experiments indicate that the ability of SON variants to restore splicing correlates with their localization to nuclear speckles, implying that proper speckle association may contribute to its regulatory activity (Fig. 5E). These observations raise the possibility that SON-dependent splicing regulation may involve both the biochemical activity of the SON protein and the spatial environment provided by nuclear speckles. Because SON depletion also alters the morphology and potentially their composition, it remains difficult to fully disentangle the contribution of SON as an individual factor from its role in nuclear speckle organization.^16^ Future studies that more precisely manipulate speckle architecture independently of SON function will be important for clarifying the relative contributions of these two components. Notably, during the preparation of this manuscript, an independent study reported that nuclear speckles are required for proper processing of RNAs derived from GC-rich isochores.^38^

## Acknowledgments

We thank members of the Yin and Li Laboratories for suggestions and insightful discussions, and S. Sun constructed the SON^dTAG^ cell line. We also wish to thank F. Chen for dTAG degron system, S. Li and the faculty of Protein Chemistry and Proteomics Platform and Metabolomics and Lipidomics Platform at Tsinghua University for mass spectrometry analysis, and W. Yin, S. Liu, Y. Huang from the Core Facilities at Zhejiang University School of Medicine for their technical support. Grant support comes from the National Natural Science Foundation of China (32270582, 32570645 and 32122019 to Y.Y.; 32171283 and 32370610 to X.L.), National Key Research and Development Program of China (2023YFA1801300 to Y.Y.), and Natural Science Foundation of Zhejiang Province, China (LZ24C060002 to Y.Y. and LR22C060002 to X.L.).

## Author contributions

Y.Y. conceived and supervised the project. X.L. provided important suggestions and help for the project. Y.Z. performed all the bioinformatics analyses. W.F., X.Z., C.T. performed most experiments. Y.Y., Y.Z., W.F., X.Z. prepared all the figures and wrote the manuscript.

## Data availability

4sU-seq, 4sU-pA-seq, TT-seq, subcellular fraction 4sU-seq, U2AF2 CLIP-seq, U2 RAP-seq, SON TurboRIP-seq, and Pol II NTD ChIP-seq data have been deposited to NCBI GEO (accession number GSE324440). All data will be publicly available as of the date of publication.

## Methods

### Cell lines, culture conditions, transfections and treatment

Mouse embryonic stem (mES) cells (WT 46C and the indicated modified derivatives) were maintained on gelatin-coated tissue plates in standard ES culture medium (DMEM; Dulbecco’s modified Eagle’s medium) supplemented with 15% heat-inactivated fetal calf serum, 0.1 mM β-mercaptoethanol, 2 mM L-glutamine, 0.1 mM non-essential amino acids, 1000 U/ml recombinant leukemia inhibitory factor (LIF) and 50 U/ml Penicillin/Streptomycin. Plasmid transfection was performed using Lipomaster 2000 (Vazyme) according to the manufacturer’s instructions. The culture medium was replaced with fresh medium containing the indicated drugs 12–24 h after transfection.

### SON^dTAG^ KI cell line construction

A homozygous SON^dTAG^ mES cell line was generated by CRISPR/Cas9-mediated homologous recombination. The FKBP12^F36V^ degron sequence was inserted in-frame at exon 2 of the endogenous *Son* locus. The donor construct comprised two homology arms (∼500 bp each) flanking a cassette encoding the FKBP12^F36V^ degron linked via a P2A peptide to a blasticidin resistance gene.

WT mESCs were co-transfected with the donor construct together with Cas9 and sgRNA expression plasmids. Following blasticidin selection, single clones were isolated and analyzed as described to confirm correct genomic integration. Clones showing efficient SON degradation upon dTAG treatment were used for subsequent experiments.

### 4sU-seq, 4sU-pA-seq, subcellular fractionation 4sU-seq, TT-seq and data analysis

SON^dTAG^ mESCs were cultured with or without dTAG for 3 h, followed by addition of 4-thiouridine (4sU; 200 μM) for 2 h to label newly synthesized RNAs. After labeling, total RNA was extracted by direct lysis in TRIzol reagent (Invitrogen). For the enrichment of nascent transcripts, 20 μg of total RNA was supplemented with 200 ng 4sU-labeled S2 RNA as an internal spike-in control and the RNA was biotinylated and enriched using streptavidin beads (Beyotime). The beads were washed four times with nuclear lysis buffer (50 mM Tris-HCl, 10 mM EDTA, 1% SDS), and bound RNA was eluted with nuclear lysis buffer containing 100 mM DTT.

For 4sU-pA-seq, 4sU-enriched RNA was further subjected to poly(A)+ selection using Dynabeads™ Oligo(dT)_25_ (Invitrogen) pretreated with 0.1 N NaOH and equilibrated in binding buffer (20 mM Tris-HCl, pH 7.5, 1 M LiCl, 2 mM EDTA). Following incubation, beads were washed with wash buffer (10 mM Tris-HCl, 0.15 M LiCl, 1 mM EDTA), and poly(A)+ RNA was eluted with 10 mM Tris-HCl at 65 °C.

For subcellular fractionation 4sU-seq, cells were labeled with 4sU as described above, and fractionation was performed as previously described.^26^ Briefly, RNA from cytoplasm, nucleoplasm, and chromatin fractions was extracted, and 4sU-labeled RNA from each fraction was subjected to biotinylation, streptavidin-mediated enrichment as described above. rRNA depletion was performed using the Ribo-off rRNA Depletion Kit (Human/Mouse/Rat; Vazyme), and RNA-seq libraries were prepared using the VAHTS Universal V10 RNA-seq Library Prep Kit for Illumina (Vazyme) according to the manufacturer’s instructions. Libraries were sequenced on an Illumina platform. For TT-seq, SON^dTAG^ mESCs were cultured with or without dTAG for 3 h, followed by 4sU (200 μM) addition for 10 min to label newly synthesized RNAs. Total RNA was extracted using TRIzol reagent (Invitrogen). The labeled RNAs were biotinylated as described in the 4sU-seq protocol and subsequently fragmented by alkaline hydrolysis (0.1× NaOH) on ice for 10 min, followed by neutralization with Tris-HCl. Fragmented RNAs were subjected to streptavidin-mediated enrichment and library preparation as described above.

All 4sU-labeled RNA-seq reads were aligned to the mouse reference genome (mm10) using Hisat2 (v2.2.1) with default parameters.^39^ Gene-level read counts were quantified with StringTie (v3.0.0; parameter: --rf) using GENCODE vM25 as the reference annotation.^40^ Differential expression analysis was performed using a two-sided *t*-test, with significance defined as *P* < 0.05.

For 3’ss usage analysis, aligned BAM files were converted to BED12 format using BEDTools (v2.31.1) and further separated into spliced and unspliced reads.^41^ These reads were intersected with annotated intronic regions. Splicing efficiency for each intron was estimated based on reads spanning the junction between the last nucleotide of the intron and the first nucleotide of the downstream exon (as illustrated in Fig. 2A). Spliced and unspliced read counts were normalized to reads per million mapped reads (RPM), and fold changes were calculated between SON^dTAG^ mESCs treated with dTAG for 5 h and untreated controls. Motif scores for 5’ss and 3’ss were computed across the mm10 genome as previously described.^42^

Alternative splicing events were identified using rMATS (v4.1.2; parameter: --cstat 0.0001).^43^ For motif analysis of exon skipping and intron retention events upon SON^dTAG^ depletion, 5,000 random 5’ss and 3’ss from the mouse genome were selected as a background control. Sequence logos were generated using WebLogo (v3.7.12) with default settings.

### Machine learning model for predicting SON-regulated introns based on intron features

To predict SON-regulated introns, we constructed a supervised machine learning framework using intron-level features. Our dataset consisted of introns classified as SON-regulated or unregulated based on SON^dTAG^ 5h 4sU-seq data. Each intron was represented by a set of features, including intron GC content and length, intron splice site strength scores, 3’ss nucleotide ratio, and speckle localization scores.

We employed a Random Forest classifier as implemented in scikit-learn. Data were split into training (80%) and test (20%) sets, stratified by class. Model performance was evaluated using ROC AUC on the held-out test set. This approach allowed identification of intron features most predictive of SON regulation.

### Pol II ChIP-seq and data analysis

WT mESCs grown in 10-cm dishes were crosslinked with 1% formaldehyde for 10 min and quenched with 0.125 M glycine. Pelleted cells were resuspended in nuclear lysis buffer supplemented with protease inhibitors and fragmented by probe sonication to ∼200-500 bp. After centrifugation, the supernatant was diluted with ChIP dilution buffer and incubated with antibody against RNAP II N-terminal (Pol II N-terminal; ABclonal) overnight at 4 °C, followed by incubation with Protein A/G magnetic beads for 3 h. Beads were washed sequentially with low-salt, high-salt, LiCl and TBST wash buffers, and bound chromatin was eluted in ChIP elution buffer. Crosslinks were reversed by proteinase K digestion at 56 °C, and DNA was purified using a PCR purification kit (Cycle Pure Kit; Omega). Libraries were prepared using standard Illumina protocols and sequenced on an Illumina platform.

Pol II ChIP-seq reads were aligned to the reference genome using Bowtie2 (v 2.5.4).^44^ The resulting BAM files were converted to BED format to facilitate downstream analysis. For each gene, the traveling ratio (TR) was calculated as the ratio of Pol II density in the promoter region (−30 to +300 relative to the TSS) to that in the gene body (the remainder of the gene).

### U2AF2 CLIP-seq and data analysis

CLIP-seq was performed as previously described with minor modifications.^24^ We co-transfected SON^dTAG^ mESCs with PB-Hygro-U2AF2-StrepII-His plasmids and pBASE. Cells were expanded in appropriate growth medium in 10-cm plates. SON^dTAG^ mESCs were cultured with or without dTAG for 2 h, followed by 4-thiouridine (4sU) addition (400 μM) for 1 h. Remove cell culture medium and cross-link the cells twice by 200-mJ 365-nm ultraviolet (UV) irradiation. Cells were lysed with CLIP lysis buffer (50 mM Tris-Cl, 100 mM NaCl and 1% NP-40), followed by digestion with RNase I and DNase I at 37 °C for 5 min to partially degrade RNA and DNA. The digestion was stopped by adding SDS to a final concentration of 0.1% and the lysate was centrifuged. Strep-Tactin®XT agarose beads (IBA Lifesciences) were added to the lysate and incubated overnight at 4 °C. Beads were washed sequentially with high-salt wash buffer (5× PBS, 0.1% SDS, 1% NP-40) and Strep wash buffer (50 mM Tris-Cl, 0.5% NP-40) supplemented with varying NaCl concentrations (500 mM, 1000 mM, 3000 mM, 300 mM). The beads were eluted with nuclear lysis buffer (50 mM Tris-Cl, 10 mM EDTA and 1% SDS). Eluents were incubated with fresh Strep-Tactin®XT beads for at least 6 h or overnight at 4 °C, and beads were washed as described above. On-bead RNA was dephosphorylated with PNK and ligated to a 3’ rApp-modified linker using RNA ligase 2 truncated KQ (New England Biolabs). RNA was eluted by treatment with proteinase K buffer (50 mM Tris-Cl, 10 mM EDTA, 1% SDS, 1 μg/μl proteinase K) at 45 °C for 1 h. Ligated RNA was purified using TRIzol reagent, and all downstream library preparation steps were performed according to the published eCLIP protocol.^45^

For U2AF2 CLIP-seq analysis, we processed the raw reads using the 4sU-seq analysis pipeline. To quantify U2AF2 enrichment at splice sites, we defined regions spanning 50 nucleotides upstream of annotated 3’ss for both regulated and unregulated introns. These regions were generated as BED intervals based on genome annotation. Read counts overlapping each region were calculated using strand-specific intersection. For each sample, read counts were normalized to RPM to account for differences in sequencing depth. To control for background signal, enrichment was first calculated relative to the corresponding input sample by dividing the RPM in CLIP samples by the RPM in input samples for each region. To assess the effect of SON depletion, enrichment values from dTAG-treated samples were further normalized to their corresponding no-dTAG controls. Specifically, the relative enrichment was calculated as the ratio of (CLIP / Input) in dTAG versus (CLIP / Input) in no-dTAG conditions. Finally, distributions of normalized enrichment were compared between regulated and unregulated introns to evaluate the impact of SON depletion on U2AF2 association at 3’ss.

### U2 RAP-seq and data analysis

The U2 RAP-seq construct was generated based on a previously reported design.^24,46^ SON^dTAG^ mESCs with or without dTAG treatment were incubated with 0.5 mg/ml 4’-aminomethyltrioxalen (AMT) on ice for 15 min, followed by irradiation at 365 nm UV for 15 min. After cell collection and washing, nuclear lysis buffer was added and samples were sonicated. Biotinylated U2 probes were hybridized at 39 °C for 1 h, followed by incubation with streptavidin beads at 39 °C for 1 h. After washing with 0.2× SSC, RNA was eluted with nuclear lysis buffer. The eluent RNAs were further reverse-crosslinked by irradiating with 254 nm UV in the presence of acridine orange for 15 min and then used for subsequent library construction.

For U2 RAP-seq and analysis, we processed the raw reads using the 4sU-seq analysis pipeline. The quantification of U2 enrichment followed the same analytical framework as used for U2AF2 CLIP. To assess the impact of SON depletion on U2 association with splice sites, we focused on regions upstream of 3’ss from both regulated and unregulated introns. U2 binding enrichment at these regions was quantified and compared between SON^dTAG^ mESCs treated with dTAG and untreated controls. This analysis was designed to evaluate differential U2 occupancy at 3’ss and to determine whether SON depletion preferentially affects U2 recruitment to regulated introns.

### TurboRIP-seq and data analysis

TurboID-tagged Son mESCs were treated with dimethyl sulfoxide (DMSO) or 50 nM PladB (10 μM stock 200×) for 2 h prior to harvesting. Following 10 min of 500 μM biotin (250 mM stock 500×) incubation, cells were cross-linked twice with 300 mJ of 254-nm ultraviolet (UV) irradiation. Pull-down was performed similarly to CLIP-seq, with the exception that only one streptavidin magnetic beads pull-down was performed. Streptavidin beads were added to the lysate and incubated for 2.5 h at 4 °C and 30 min at room temperature. Beads were washed twice with stringent wash buffer I (50 mM Tris-Cl and 2% SDS), twice with stringent wash buffer II (5× PBS, 0.5% NP-40, 0.5% SDS and 1 M urea) and once with nuclear lysis buffer. TurboRIP was performed with mild RNase I treatment to maintain RNA integrity. Streptavidin bead pull-down was performed, followed by Proteinase K digestion to release bound RNA. The extracted RNA was subsequently used for library preparation following the RNA-seq protocol described above.

Raw reads from TurboRIP-seq were processed using the same pipeline as described for 4sU-seq data. To investigate the effect of splicing inhibition on SON association, we quantified gene-level enrichment by calculating the ratio of FPKM values in TurboRIP samples relative to their corresponding input controls. Comparisons were performed between cells treated with PladB and DMSO-treated controls. Genes were stratified into regulated and unregulated groups based on intron regulation status. The regulated gene set was defined as the union of host genes containing regulated introns identified from both SON^dTAG^ 4sU-seq and 4sU-pA-seq analyses. For each group, changes in SON association upon PladB treatment were assessed by comparing enrichment values between PladB-treated and control conditions. This analysis was designed to determine whether splicing inhibition differentially affects SON binding dynamics between regulated and unregulated gene sets.

### TurboID and mass spectrometry analysis

The TurboID protocol was modified from a previous report.^24^ Cell preparation work is the same as TurboRIP. PladB-treated and untreated SON^TurboID^ mES cells were cultured with 500 μM biotin for 1 h. Nuclei were isolated by suspending the cells with hypotonic buffer (20 mM HEPES, 10 mM KCl, 1.5 mM MgCl_2_, 1 mM EDTA, 0.1 mM Na_3_VO_4_, 0.1% NP-40 and 10% glycerol), and the isolated nuclei were resuspended in IP150 buffer supplemented with Benzonase nuclease. After incubation and centrifugation, streptavidin agarose beads were added to the supernatant. The mixture was incubated with gentle rotation at 4 °C for 2 h, followed by incubation at room temperature for 30 min. Beads were washed with stringent wash buffers following the TurboRIP protocol mentioned above. Additionally, the beads were washed once with nuclear lysis buffer and eluted twice with nuclear lysis buffer supplemented with 2 mM biotin and 20 mM dithiothreitol (DTT) at 95 °C for 5 min per elution. The combined eluates were subjected to western blot analysis and mass spectrometry.

### Immunofluorescence, microscopic imaging and analysis

The immunofluorescence of SC35 (SON^dTAG^ mESCs) was modified from a previous report.^47^ Briefly, SON^dTAG^ mESCs were seeded on Matrigel-coated coverslips. The cells were sequentially crosslinked with ethanol and 4% paraformaldehyde (PFA), then permeabilized and blocked in PBS with 5% BSA and 0.3% Triton X-100. Cells were incubated with a primary antibody against SC35 for 2 h, washed, and incubated with fluorescent secondary antibodies. After nuclear staining with DAPI, the cells were mounted for imaging.

All microscopy images were captured using an LSM Zeiss 900 inverted confocal laser scanning microscope (Axio Observer.Z1/7) with Plan-Apochromat ×63/1.40 oil objectives, using Airyscan with a GaAsp-PMT detector in super-resolution mode. Single optical sections at the maximal nuclear cross-section were acquired to visualize puncta formation. Fluorescence intensity profiles were obtained using ZEN 2.3 lite and reanalyzed in GraphPad. Condensate numbers and areas were analyzed with Fiji (v1.53c).

### Truncation rescue assay

SON truncation constructs were generated in the PiggyBac backbone. Plasmids were cotransfected with pBASE into SON^dTAG^ mESCs, followed by drug selection for 3 days. For splicing analysis, cells were treated with dTAG for 4 h before RNA extraction and RT–qPCR analysis. For competition assays, cells expressing the truncation constructs were mixed with 20% GFP-labeled *Oct4* mESCs and cultured in the presence of dTAG. The fraction of GFP-negative cells was measured by fluorescence-activated cell sorting (FACS) at the indicated time points. Rescue efficiency was normalized to cells expressing full-length (FL) SON from the same transfection batch. Cells expressing empty vector (EV) served as a negative control. The flow cytometry gating strategy was as follows: FSC/SSC parameters were used to exclude debris, and FSC-A/FSC-H were used to discriminate single cells from doublets. Samples without GFP reporter transfection were used to establish boundaries between negative and positive cells.

### Short intron definition across different species

Our analytical framework was adapted from a previously described approach.^4^ To systematically characterize intron length features across evolution, we collected genome annotations from representative species spanning lower to higher eukaryotes.

For each species, intron coordinates were extracted from gene annotation files, and intron lengths were calculated genome-wide. Introns were then classified into length categories using a percentile-based approach. Specifically, introns with lengths below the 25th percentile of the genome-wide intron length distribution were defined as “short” introns, whereas those above the 75th percentile were defined as “long” introns. This normalization strategy accounts for differences in genome architecture and intron length distributions across species. For exon-centered analyses, exons were categorized based on the lengths of their flanking introns. An exon was defined as flanked by “short” introns only if both its upstream and downstream introns were shorter than the 25th percentile threshold. Conversely, exons were classified as flanked by “long” introns when both adjacent introns exceeded the 75th percentile threshold. Exons that did not meet either criterion were excluded from further analysis to ensure clear classification.

To further investigate sequence features associated with intron length, GC content was calculated for short introns across species as the proportion of G and C nucleotides within each sequence. Comparative analyses were performed to evaluate evolutionary trends in GC composition among short introns.

### Other analysis

For gene ontology (GO) analysis, we searched for significantly enriched GO categories using DAVID Functional Annotation Bioinformatics Microarray Analysis tool (https://david.ncifcrf.gov/).^48^ Nuclear speckle scores were derived following the method reported in Bhat et al.^13^ Branchpoint sites were identified using the LaBranchoR software,^49^ which predicts branchpoint locations based on sequence features. The species phylogenetic tree was generated using the TimeTree database (https://www.timetree.org). The intrinsically disordered regions (IDRs) of the SON protein were predicted using the online tool IUPred3 (https://iupred3.elte.hu/). Representative protein sequences for each species were downloaded from NCBI or UniProt database, with the most canonical isoform selected. Sequences were aligned using the MUSCLE algorithm in MEGA (v12), and the alignment was visualized with Jalview (v2.11.5.1).

### Statistical analysis

Tables and graphs for statistical analysis were created using GraphPad Prism 8. Statistical significance between two groups was determined by a two-sided Student’s *t-*test, two-way ANOVA or Fisher’s exact test, with details indicated in figure legends. For all statistics, exact *n* values and *P* values are provided in the corresponding figure legends. All data in the graphs are shown as mean ± s.d. unless stated otherwise. Most figures and graphical representations in this study were generated using the R software environment (v4.4.1).

---

**Figure S1.**
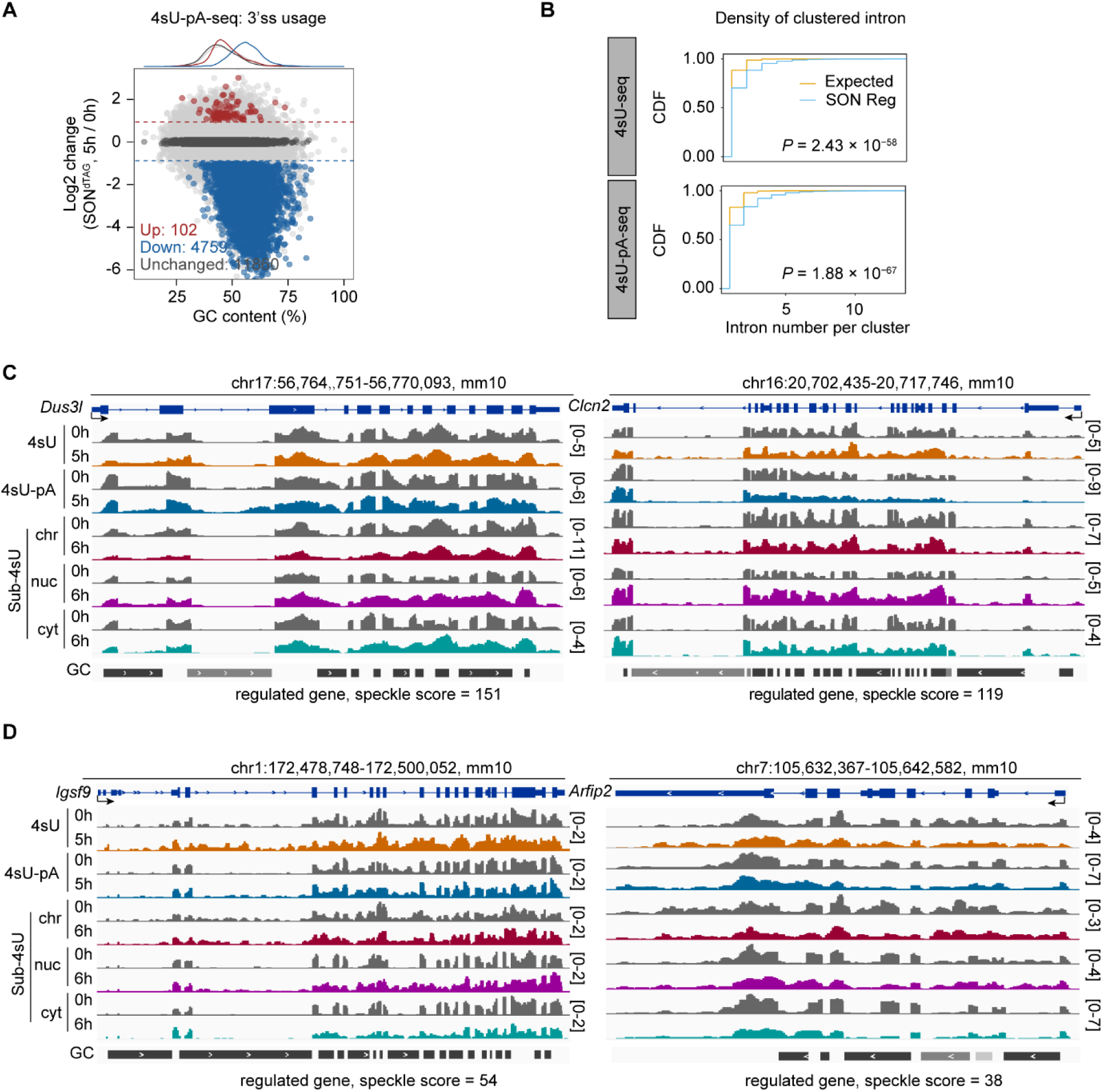
SON-regulated introns exhibit spatial clustering and are not strictly dependent on nuclear speckle proximity. (A) 3’ss usage analysis in SON^dTAG^ mESCs by 4sU-pA-seq reveals that SON-regulated introns have high GC content. (B) Comparison of intron number per cluster between expected and SON-regulated introns, with both 4sU-seq and 4sU-pA-seq datasets showing a higher proportion of SON-regulated introns within clusters. (C) IGV tracks showing two nuclear speckle–proximal genes, *Dus3l* and *Clcn2*, as SON-regulated genes. (D) IGV tracks showing two nuclear speckle–distal genes, *Igsf9* and *Arfip2*, as SON-regulated genes.

**Figure S2.**
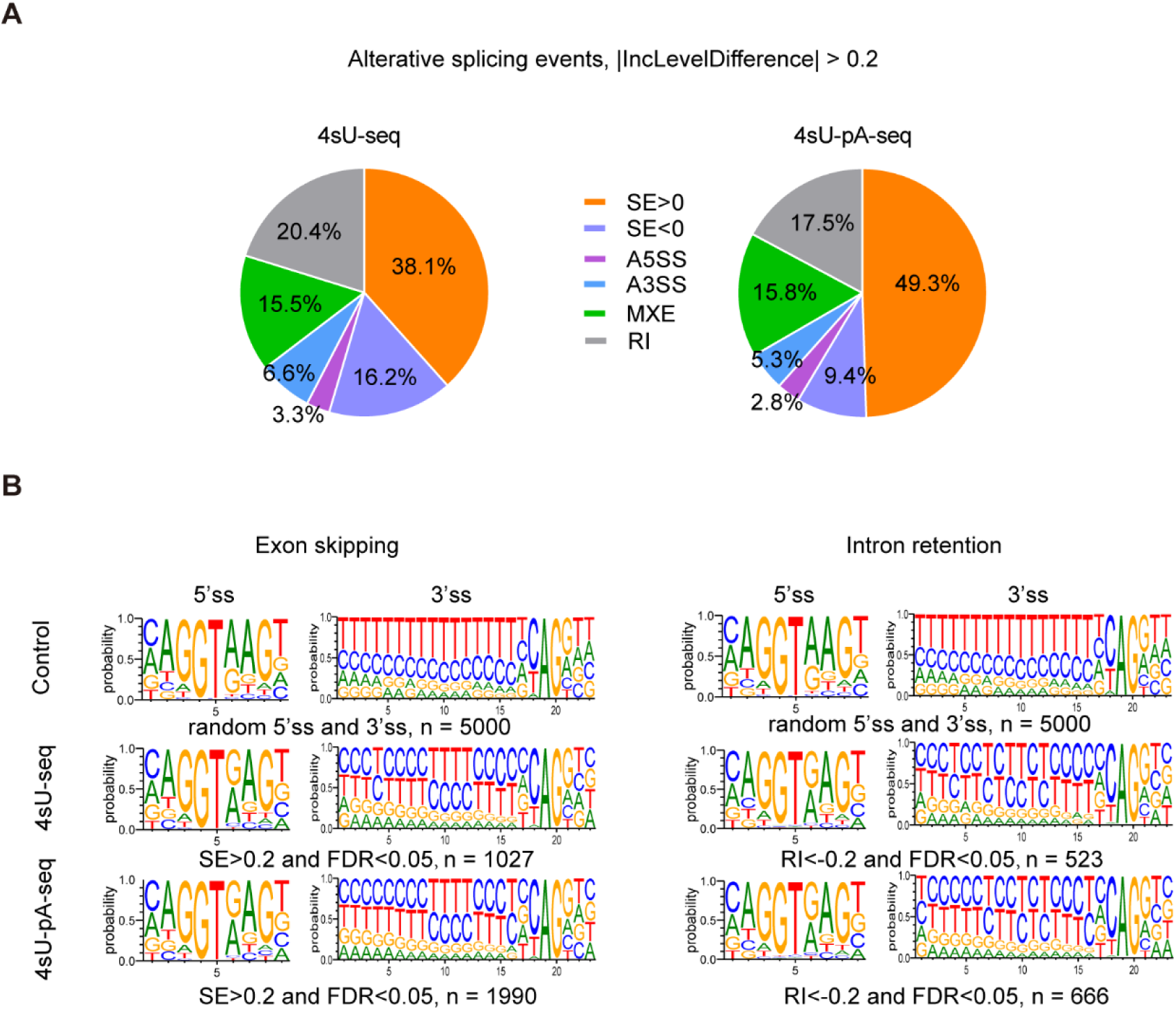
SON depletion–induced alternative splicing preferentially occurs at introns with C-rich, weak 3’ss. (A) Pie charts showing the distribution of alternative splicing events identified by rMATS in 4sU-seq (*n =* 2,695) and 4sU-pA-seq (*n =* 4,033) datasets (|IncLevelDifference| > 0.2, FDR < 0.05). (B) Comparative analysis of splice site features for regulated alternative splicing events. Left: 5’ss and 3’ss motifs of skipped exon (SE) events (IncLevelDifference > 0.2, FDR < 0.05) in control (5,000 randomly selected genomic exons), 4sU-seq (*n =* 1,027), and 4sU-pA-seq (*n =* 1,990) datasets. Right: 5’ss and 3’ss motifs of retained intron (RI) events (IncLevelDifference < −0.2, FDR < 0.05; 4sU-seq: 523; 4sU-pA-seq: 666) across the same groups. SON-regulated events exhibit minimal differences at 5’ss but show C-rich features at 3’ss compared to controls.

**Figure S3.**
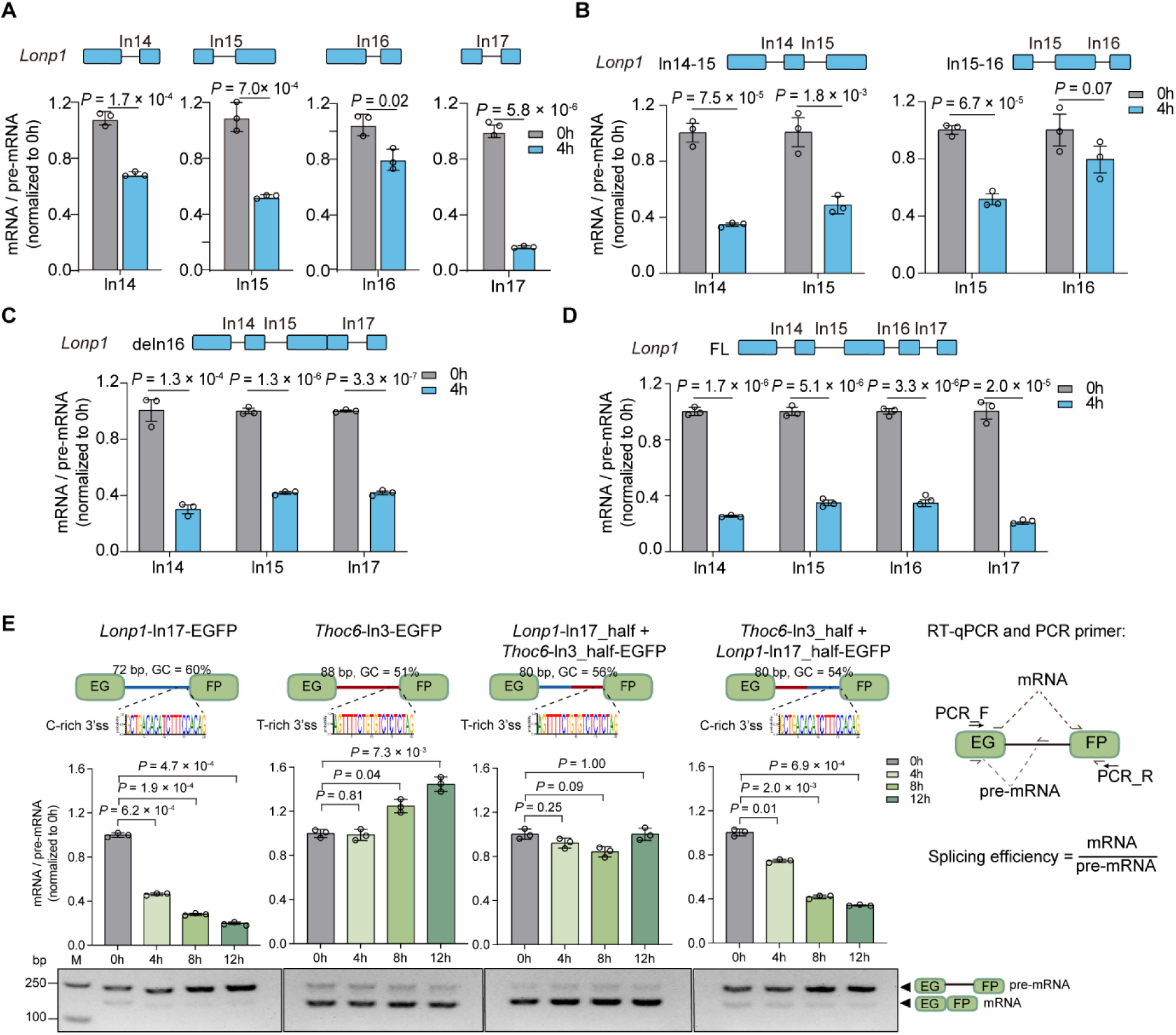
Reporter validation of SON-regulated intron splicing. (A–D) RT–qPCR analysis of splicing efficiency (mRNA/pre-mRNA) for *Lonp1* reporter constructs in SON^dTAG^ mESCs at 0 h and 4 h after dTAG treatment. Schematics (top) depict reporter transcripts with exons (blue boxes) and assayed introns (black). (A) Single-intron reporters (*Lonp1*-In14, -In15, -In16, -In17). (B) Dual-intron reporters (*Lonp1*-In14-In15 and *Lonp1*-In15-In16), containing the indicated adjacent introns from the *Lonp1* gene. (C) Intron 16 deletion reporter (*Lonp1*-deIn16). (D) Full-length reporter (*Lonp1*-FL). (E) Top, schematics of EGFP minigene reporters containing an EGFP coding sequence interrupted by full-length or chimeric introns, including *Lonp1* intron 17 (*Lonp1*-In17-EGFP), Thoc6 intron 3 (Thoc6-In3-EGFP), and chimeric constructs combining the 5’ and 3’ halves of these introns. The sequence features of the 3’ss are indicated, along with primer binding sites (EG, 5’ EGFP; FP, 3’ EGFP) used to distinguish pre-mRNA and mRNA. Bottom, RT–qPCR analysis of splicing efficiency at 0, 4, 8, and 12 h after dTAG treatment, and RT–PCR analysis of splicing products. For all panels, splicing efficiency is shown as relative mRNA/pre-mRNA levels normalized to *Actb*-E (A–D) or EGFP exon (EGFP-E) (E). Data are presented as mean ± s.d. (*n =* 3 biological replicates); *P* value*s*, two-sided *t*-test.

**Figure S4.**
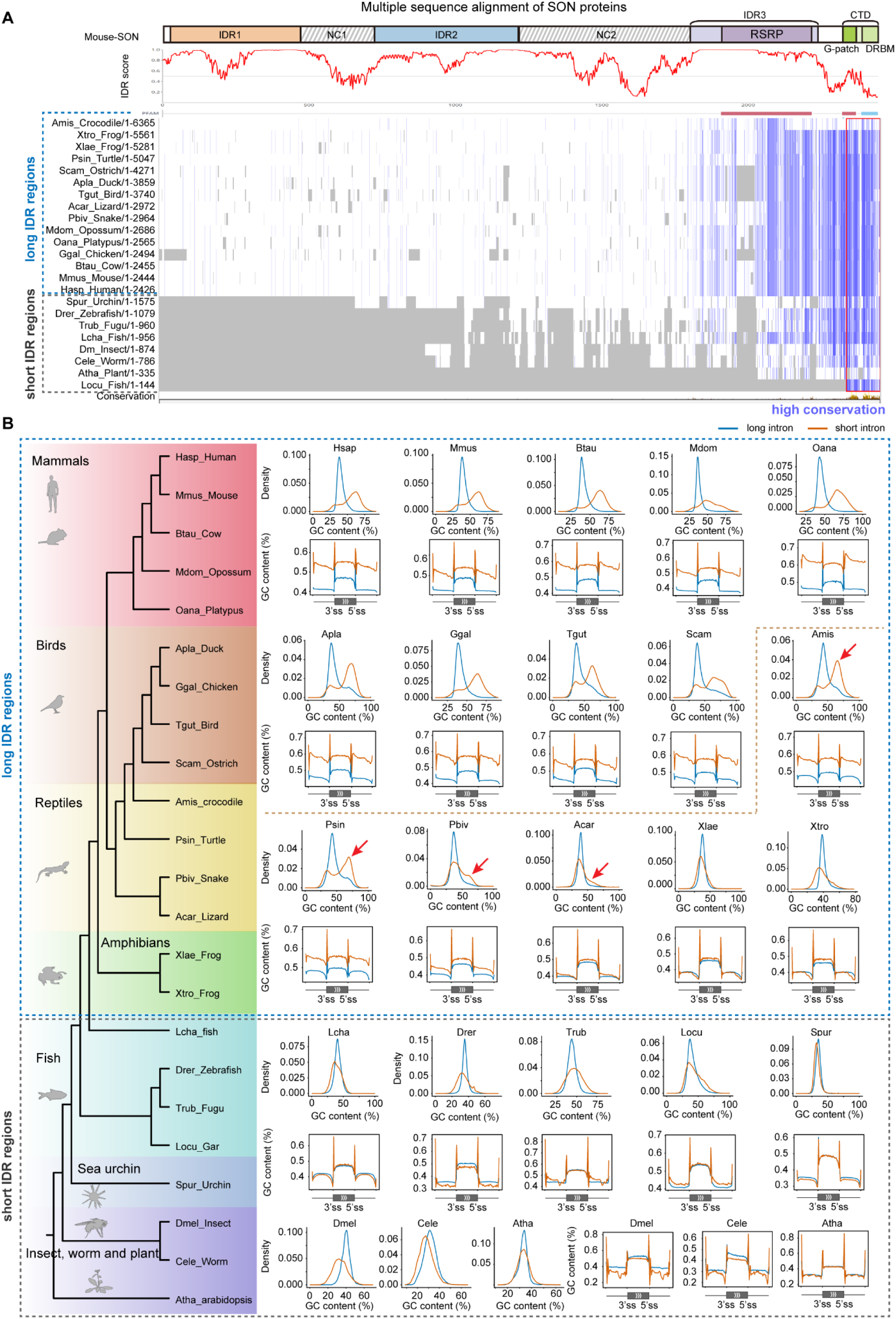
Evolutionary emergence of GC-rich short introns correlates with expansion of the SON’s IDR region. (A) Evolutionary expansion of the N-terminal IDR of SON. Top, domain architecture and IDR prediction of mouse SON along the protein sequence (predicted by IUPred3). Bottom, multiple sequence alignment of SON orthologs from representative species, showing conservation patterns across evolution. Each row represents one species, and color intensity indicates sequence conservation. Gray indicates missing sequence. Vertebrate SON proteins exhibit an expanded N-terminal disordered region, whereas the C-terminal region containing conserved domains remains highly conserved. (B) Species are arranged according to a phylogenetic tree generated using TimeTree5 and grouped into major clades (mammals, birds, reptiles, amphibians, fish, and invertebrates/plant). Based on SON’s IDR length, species were grouped into two groups indicated by dashed boxes: species with expanded IDRs (upper panel) and species with short IDRs (lower panel). For each species, two analyses are shown. The top density plot shows the GC content distribution of long introns and short introns, illustrating the emergence of GC-rich introns during evolution. The bottom plot shows the average GC content along intron regions aligned at splice sites. Introns and exons shorter than 50 nt were excluded; short exons and their flanking introns were divided into 50 bins, and the average GC content in each bin was calculated and plotted. Species with short SON’s IDRs show relatively uniform intron GC content, whereas species with expanded SON’s IDRs display introns with elevated GC levels. Red arrows indicate representative species in which GC-rich short introns are observed. This feature becomes evident in amphibians and reptiles and is further enhanced in birds and mammals, coinciding with the evolutionary expansion of the SON N-terminal IDR.

## References

1 Bernardi, G. The Isochore Organization of the Human Genome and Its Evolutionary History - a Review. Gene 135, 57–66, doi:Doi 10.1016/0378-1119(93)90049-9 (1993).

2 Lander, E. S. et al. Initial sequencing and analysis of the human genome. Nature 409, 860–921, doi:10.1038/35057062 (2001).

3 Birney, E. et al. Identification and analysis of functional elements in 1% of the human genome by the ENCODE pilot project. Nature 447, 799–816, doi:10.1038/nature05874 (2007).

4 Amit, M. et al. Differential GC content between exons and introns establishes distinct strategies of splice-site recognition. Cell Rep 1, 543–556, doi:10.1016/j.celrep.2012.03.013 (2012).

5 Tammer, L. et al. Gene architecture directs splicing outcome in separate nuclear spatial regions. Mol Cell 82, 1021–1034.e1028, doi:10.1016/j.molcel.2022.02.001 (2022).

6 Fu, X. D. & Ares, M., Jr. Context-dependent control of alternative splicing by RNA-binding proteins. Nat Rev Genet 15, 689–701, doi:10.1038/nrg3778 (2014).

7 Will, C. L. & Luhrmann, R. Spliceosome structure and function. Cold Spring Harb Perspect Biol 3, doi:10.1101/cshperspect.a003707 (2011).

8 Chen, Y. et al. Mapping 3D genome organization relative to nuclear compartments using TSA-Seq as a cytological ruler. J Cell Biol 217, 4025–4048, doi:10.1083/jcb.201807108 (2018).

9 Spector, D. L. & Lamond, A. I. Nuclear speckles. Cold Spring Harb Perspect Biol 3, doi:10.1101/cshperspect.a000646 (2011).

10 Faber, G. P., Nadav-Eliyahu, S. & Shav-Tal, Y. Nuclear speckles – a driving force in gene expression. J Cell Sci 135, doi:10.1242/jcs.259594 (2022).

11 Ilık, İ. A. & Aktaş, T. Nuclear speckles: dynamic hubs of gene expression regulation. The FEBS Journal 289, 7234–7245, doi:10.1111/febs.16117 (2021).

12 Fan, J. et al. mRNAs are sorted for export or degradation before passing through nuclear speckles. Nucleic Acids Res 46, 8404–8416, doi:10.1093/nar/gky650 (2018).

13 Bhat, P. et al. Genome organization around nuclear speckles drives mRNA splicing efficiency. Nature 629, 1165–1173, doi:10.1038/s41586-024-07429-6 (2024).

14 Wu, J. et al. Dynamics of RNA localization to nuclear speckles are connected to splicing efficiency. Sci Adv 10, eadp7727, doi:10.1126/sciadv.adp7727 (2024).

15 Zhang, L. G. et al. TSA-seq reveals a largely conserved genome organization relative to nuclear speckles with small position changes tightly correlated with gene expression changes. Genome Res 31, 251–264, doi:10.1101/gr.266239.120 (2021).

16 Ilik, I. A. et al. SON and SRRM2 are essential for nuclear speckle formation. Elife 9, doi:10.7554/eLife.60579 (2020).

17 Xu, S. et al. SRRM2 organizes splicing condensates to regulate alternative splicing. Nucleic Acids Res 50, 8599–8614, doi:10.1093/nar/gkac669 (2022).

18 Ahn, E. Y. et al. SON controls cell-cycle progression by coordinated regulation of RNA splicing. Mol Cell 42, 185–198, doi:10.1016/j.molcel.2011.03.014 (2011).

19 Lu, X. et al. SON connects the splicing-regulatory network with pluripotency in human embryonic stem cells. Nat Cell Biol 15, 1141–1152, doi:10.1038/ncb2839 (2013).

20 Zhu, X. L. et al. Whole-exome sequencing in undiagnosed genetic diseases: interpreting 119 trios. Genet Med 17, 774–781, doi:10.1038/gim.2014.191 (2015).

21 Tokita, M. J. et al. De Novo Truncating Variants in SON Cause Intellectual Disability, Congenital Malformations, and Failure to Thrive. Am J Hum Genet 99, 720–727, doi:10.1016/j.ajhg.2016.06.035 (2016).

22 Kim, J. H. et al. De Novo Mutations in SON Disrupt RNA Splicing of Genes Essential for Brain Development and Metabolism, Causing an Intellectual-Disability Syndrome. Am J Hum Genet 99, 711–719, doi:10.1016/j.ajhg.2016.06.029 (2016).

23 Nabet, B. et al. The dTAG system for immediate and target-specific protein degradation. Nat Chem Biol 14, 431–441, doi:10.1038/s41589-018-0021-8 (2018).

24 Zhou, Y. et al. Condensation of ZFP207 and U1 snRNP promotes spliceosome assembly. Nat Struct Mol Biol 32, 1038–1049, doi:10.1038/s41594-025-01501-z (2025).

25 Schwalb, B. et al. TT-seq maps the human transient transcriptome. Science 352, 1225–1228, doi:10.1126/science.aad9841 (2016).

26 Bhatt, D. M. et al. Transcript dynamics of proinflammatory genes revealed by sequence analysis of subcellular RNA fractions. Cell 150, 279–290, doi:10.1016/j.cell.2012.05.043 (2012).

27 Reppas, N. B., Wade, J. T., Church, G. M. & Struhl, K. The transition between transcriptional initiation and elongation in E. coli is highly variable and often rate limiting. Mol Cell 24, 747–757, doi:10.1016/j.molcel.2006.10.030 (2006).

28 Roscigno, R. F., Weiner, M. & Garciablanco, M. A. A Mutational Analysis of the Polypyrimidine Tract of Introns - Effects of Sequence Differences in Pyrimidine Tracts on Splicing. J Biol Chem 268, 11222–11229 (1993).

29 Glasser, E. et al. Pre-mRNA splicing factor U2AF2 recognizes distinct conformations of nucleotide variants at the center of the pre-mRNA splice site signal. Nucleic Acids Res 50, 5299–5312, doi:10.1093/nar/gkac287 (2022).

30 Cho, K. F. et al. Proximity labeling in mammalian cells with TurboID and split-TurboID. Nat Protoc 15, 3971–3999, doi:10.1038/s41596-020-0399-0 (2020).

31 Kotake, Y. et al. Splicing factor SF3b as a target of the antitumor natural product pladienolide. Nat Chem Biol 3, 570–575, doi:10.1038/nchembio.2007.16 (2007).

32 Rahl, P. B. et al. c-Myc regulates transcriptional pause release. Cell 141, 432–445, doi:10.1016/j.cell.2010.03.030 (2010).

33 Zamft, B., Bintu, L., Ishibashi, T. & Bustamante, C. Nascent RNA structure modulates the transcriptional dynamics of RNA polymerases. Proc Natl Acad Sci U S A 109, 8948–8953, doi:10.1073/pnas.1205063109 (2012).

34 Jonkers, I., Kwak, H. & Lis, J. T. Genome-wide dynamics of Pol II elongation and its interplay with promoter proximal pausing, chromatin, and exons. Elife 3, doi:10.7554/eLife.02407 (2014).

35 Veloso, A. et al. Rate of elongation by RNA polymerase II is associated with specific gene features and epigenetic modifications. Genome Res 24, 896–905, doi:10.1101/gr.171405.113 (2014).

36 Wegener, M. & Müller-McNicoll, M. View from an mRNP: The Roles of SR Proteins in Assembly, Maturation and Turnover. Biology of Mrna: Structure and Function 1203, 83–112, doi:10.1007/978-3-030-31434-7_3 (2019).

37 Pandit, S. et al. Genome-wide Analysis Reveals SR Protein Cooperation and Competition in Regulated Splicing. Molecular Cell 50, 223–235, doi:10.1016/j.molcel.2013.03.001 (2013).

38 Małszycki, M. et al. Nuclear speckles enable processing of RNA from GC-rich isochores. Cell, doi:10.1016/j.cell.2026.01.011 (2026).

39 Kim, D., Paggi, J. M., Park, C., Bennett, C. & Salzberg, S. L. Graph-based genome alignment and genotyping with HISAT2 and HISAT-genotype. Nat Biotechnol 37, 907–915, doi:10.1038/s41587-019-0201-4 (2019).

40 Kovaka, S. et al. Transcriptome assembly from long-read RNA-seq alignments with StringTie2. Genome Biol 20, 278, doi:10.1186/s13059-019-1910-1 (2019).

41 Quinlan, A. R. & Hall, I. M. BEDTools: a flexible suite of utilities for comparing genomic features. Bioinformatics 26, 841–842, doi:10.1093/bioinformatics/btq033 (2010).

42 Almada, A. E., Wu, X. B., Kriz, A. J., Burge, C. B. & Sharp, P. A. Promoter directionality is controlled by U1 snRNP and polyadenylation signals. Nature 499, 360–U141, doi:10.1038/nature12349 (2013).

43 Shen, S. et al. rMATS: robust and flexible detection of differential alternative splicing from replicate RNA-Seq data. Proc Natl Acad Sci U S A 111, E5593–5601, doi:10.1073/pnas.1419161111 (2014).

44 Langmead, B. & Salzberg, S. L. Fast gapped-read alignment with Bowtie 2. Nat Methods 9, 357–U354, doi:10.1038/Nmeth.1923 (2012).

45 Van Nostrand, E. L. et al. Robust transcriptome-wide discovery of RNA-binding protein binding sites with enhanced CLIP (eCLIP). Nat Methods 13, 508–514, doi:10.1038/nmeth.3810 (2016).

46 Engreitz, J. M. et al. RNA-RNA interactions enable specific targeting of noncoding RNAs to nascent Pre-mRNAs and chromatin sites. Cell 159, 188–199, doi:10.1016/j.cell.2014.08.018 (2014).

47 Zhou, Y. et al. Condensation of ZFP207 and U1 snRNP promotes spliceosome assembly. Nature Structural & Molecular Biology 32, 1038–1049, doi:10.1038/s41594-025-01501-z (2025).

48 Sherman, B. T. et al. DAVID: a web server for functional enrichment analysis and functional annotation of gene lists (2021 update). Nucleic Acids Res 50, W216–W221, doi:10.1093/nar/gkac194 (2022).

49 Paggi, J. M. & Bejerano, G. A sequence-based, deep learning model accurately predicts RNA splicing branchpoints. RNA 24, 1647–1658, doi:10.1261/rna.066290.118 (2018).

